# Association of the benzoxazinoid pathway with boron homeostasis in maize

**DOI:** 10.1101/2024.10.25.620361

**Authors:** Liuyang Chu, Vivek Shrestha, Cay Christin Schäfer, Jan Niedens, George W. Meyer, Zoe Darnell, Tyler Kling, Tobias Dürr-Mayer, Aleksej Abramov, Monika Frey, Henning Jessen, Gabriel Schaaf, Frank Hochholdinger, Agnieszka Nowak-Król, Paula McSteen, Ruthie Angelovici, Michaela S. Matthes

## Abstract

Both deficiency and toxicity of the micronutrient boron lead to severe reductions in crop yield. Despite this agricultural importance, the molecular basis underlying boron homeostasis in plants is not resolved. To identify molecular players involved in boron homeostasis in maize (*Zea mays*), we measured boron levels in the Goodman-Buckler association panel and performed genome-wide association studies. These analyses detected the *benzoxazinless* (*bx*) gene, *bx3*, involved in the biosynthesis of benzoxazinoids, like DIMBOA, major defense compounds in maize. Genes involved in DIMBOA biosynthesis are all located in close proximity in the genome and the biosynthesis mutants, including *bx3* are all DIMBOA deficient. We show that the *bx3* mutant has enhanced boron concentration in leaves compared to the B73 control plants, which correlates with enhanced leaf tip necrosis, a phenotype associated with boron toxicity. In contrast, other DIMBOA-deficient maize mutants did not show altered boron levels nor the leaf tip necrosis phenotype, suggesting that boron is not associated with DIMBOA. Instead, our analyses suggest that the accumulation of boron is linked to the benzoxazinoid intermediates indolin-2-one (ION) and 3-hydroxy-ION. Therefore, our results connect boron homeostasis to the benzoxazinoid plant defense pathway through *bx3* and specific intermediates, rendering the benzoxazinoid biosynthesis pathway a potential target for crop improvement in inadequate boron conditions.

**One sentence summary:** GWAS identified *benzoxazinless3* in the benzoxazinoid biosynthesis pathway to play a role in boron homeostasis in maize.

## Introduction

The micronutrient boron is essential for proper plant growth (Warington, 1923) and both deficiencies and toxicities of boron in the soil lead to severe yield reductions in many major crops, including maize (Brdar-Jokanović, 2020). Soils with either low or toxic boron levels are widespread in the world (Landi et al., 2019; Brdar-Jokanović, 2020 and references therein), making the study of the effects of boron supply on plant development an important topic for agriculture. After uptake from the soil, boron moves along the transpiration stream and accumulates at its end. Consequently, excess boron accumulates at the leaf blade tips and extends into the margins (Brown and Shelp, 1997; Marschner, 2012), which causes leaf necrosis when boron supply is too high (Eaton, 1944; Reid et al., 2004). Boron toxicity leads to reduced chlorophyll content, stomatal conductance, and photosynthesis (as reviewed in Landi et al., 2019). Boron deficiency primarily affects meristematic tissues (Sommer and Sorokin, 1928), leading to reductions in leaf blade width and length, sterility, and in severe cases to seedling lethality. In the root, both boron deficiency and toxicity lead to a reduction of primary root length (Reid et al., 2004; Choi et al., 2007; Aquea et al., 2012; Esim et al., 2013; Poza-Viejo et al., 2018; Zhang et al., 2020), and to a reduction in lateral root density (Housh et al., 2020; Wilder et al., 2022). Understanding the molecular mechanisms that allow for efficiently using low levels or tolerating toxic levels of boron, will help in developing efficient and tolerant varieties providing a sustainable solution to overcome yield reductions due to suboptimal or toxic boron supply.

The main characterized function of boron is in crosslinking of the pectin subunits rhamnogalacturonan-II (RG-II) in the cell wall (Kobayashi et al., 1996; Matoh et al., 1996; O’Neill et al., 1996) and boron efficiency processes include boron uptake, boron trans-and retranslocation, and boron utilization. From these processes, genetic components involved in boron uptake are currently best understood in the model organism *Arabidopsis thaliana* (Arabidopsis). Boron is taken up by plants in the form of boric acid and transported within plants in the form of the borate anion (for a recent review see Onuh and Miwa, 2021). While passive uptake prevails in boron adequate soil conditions, facilitated and active boron transport is needed in low and excess soil boron conditions. Boron transporters belong to the nodulin 26-like intrinsic protein (NIP) subfamily of the major intrinsic protein (MIP) family of boric acid importers and the BOR family of borate exporters. In Arabidopsis, the three NIPs, involved in boron import, and the seven BORs, involved in boron export, have tissue and functional specificities (as reviewed in Onuh and Miwa, 2021). The major players are *At*NIP5;1 (Takano et al., 2006) and *At*BOR1 (Noguchi et al., 1997). Orthologs of these genes have been characterized in many plant species, including maize (Chatterjee et al., 2014; Durbak et al., 2014; Leonard et al., 2014; Chatterjee et al., 2017). *Zmtassel-less1* (*Zmtls1)*, is the *AtNIP5;1* co-ortholog in maize and shows seedling lethality, when grown in low boron conditions. When grown in adequate boron conditions, only reproductive defects are prominent (Durbak et al., 2014; Leonard et al., 2014). Due to impaired boron uptake, this mutant is inherently boron deficient in meristem tissues and all defects can be rescued by boron fertilization (Durbak et al., 2014; Matthes et al., 2018), making it a good tool to study boron deficiency responses in maize (Matthes et al., 2022; Matthes et al., 2023). Boron transporters additionally were shown to provide molecular targets for engineering plants adapted to altered soil boron concentrations (Miwa et al., 2006; Sutton et al., 2007; Kato et al., 2009; Schnurbusch et al., 2010; Uraguchi et al., 2014; Hayes et al., 2015; He et al., 2021).

Several lines of evidence indicate that additional molecular candidates exist that either regulate intracellular levels of boron or adapt cellular metabolism to changing boron levels. For example, various classes of genes are differentially regulated in boron deficient and toxic conditions (Peng et al., 2012) and there are striking differences between phenotypes of boron transporter mutants and phenotypes of plants grown in boron deficient conditions (as reviewed in Matthes et al., 2020). Numerous studies additionally highlight the importance of meristem genes and phytohormone cascades in the adaptation of plants to low boron levels (Abreu et al., 2014; Eggert and von Wirén, 2017; Poza-Viejo et al., 2018; Gómez-Soto et al., 2019; Matthes et al., 2022; Pommerrenig et al., 2022; Matthes et al., 2023). Recently the boron deficiency response was additionally shown to reflect a wounding response in *Brassica napus* (Verwaaijen et al., 2023). These examples suggest that there are additional pathways that regulate either plant boron levels or susceptibilities and therefore might be harnessed for developing high yielding crops in boron deficient conditions.

In Arabidopsis, few non-boron transporter genes potentially involved in boron homeostasis were identified. Of those, specific transcription factors were identified that regulate boron import by directly binding to the promoter of boron transporter gene *NIP5;1* (Kasajima and Fujiwara, 2007; Kasajima et al., 2010; Feng et al., 2020; Zhang et al., 2024). For others, a link to boron transport remains elusive and they might therefore act independent of it. These include, the *low boron tolerance 1* mutant (Huai et al., 2018), for which the underlying gene remains to be identified, the *sensitive to high-level of boron 1* mutant, which is defective in *At*HY1, involved in photomorphogenesis (Lv et al., 2017), genes encoding specific subunits of condensin II (Sakamoto et al., 2011), affecting RG-II biosynthesis or dimer formation (O’Neill et al., 2001; Sechet et al., 2018) or genes influencing the deposition of pectin in the cell wall (Hiroguchi et al., 2021). More recently evidence from Arabidopsis and *Rosa* cell suspension cultures emphasized the importance of a properly functioning Golgi apparatus for boron related processes (Chormova et al., 2014; Chormova and Fry, 2016; Begum and Fry, 2022; Onuh and Miwa, 2023).

Efforts to elucidate molecular mechanisms underlying boron homeostasis in crops are ongoing and include the assessment of genetic variation (Pommerrenig et al., 2018; He et al., 2021), QTL analyses (Paull et al., 1991; Jefferies et al., 1999; Jefferies et al., 2000; Xu et al., 2001; Sutton et al., 2007; Schnurbusch et al., 2010; Zhao et al., 2012; Pallotta et al., 2014) or GWAS (de Abreu Neto et al., 2017; Jia et al., 2021). Furthermore, transcriptome studies identified genes that respond to varying boron levels in the soil or growing media (Zeng et al., 2008). While such efforts detected various genomic regions and a multitude of genes affected by altered soil boron levels, the elucidation of molecular players instrumental for boron homeostasis, and therefore deficiency or toxicity tolerance, is not resolved.

Here, we report the detection of the benzoxazinoid biosynthesis gene *benzoxazinless3* (*bx3*) as a molecular candidate associated with boron homeostasis in the major crop maize (*Zea mays* L.). Our study links altered boron levels with *bx3* and the benzoxazinoid pathway intermediates indolin-2-one and 3-hydroxyindolin-2-one. Benzoxazinoids are indole-derived secondary defense metabolites, which have best been studied in grasses (Poacea) and additionally independently evolved in some eudicot species (Florean et al., 2023). They not only regulate below and above-ground biotic interactions but are also known as signaling molecules and as iron chelators supporting iron uptake (Hu et al., 2018). Our study therefore adds another layer to the multifunctionality of benzoxazinoids and suggests the benzoxazinoid pathway as target for engineering plants adapted to altered soil boron concentrations.

## Material and Methods

### Germplasm and plant growing conditions

277 inbred lines of the 282 Goodman-Buckler association panel (Flint-Garcia et al., 2005), consisting of diverse maize lines including tropical, sub-tropical, temperate, sweet corn, and popcorn lines, were grown in a randomized complete block design in the summers of 2017 and 2018 at Genetics Farm of the University of Missouri (Columbia, Missouri, USA). Soil boron concentration in both years was 0.39 mg/kg as evaluated with the azomethine H method (Lohse, 1982) by the Soil and Plant Testing Laboratory of the University of Missouri, which can be considered low considering the recommendation to apply boron to field maize, when soil levels are < 0.75 ppm (Heckmann, 2009).

The alleles used for the respective mutants were *Bx3::Mu* (*bx3*) (Frey et al., 1997), *bx1::Mu* (*bx1*) (Frey et al., 1997), *bx2::Ds* (*bx2*) (Tzin et al., 2017), and *tls1-ref* (*tls1*) (Durbak et al., 2014). The *bx1*, *bx3*, and *tls1* mutants were backcrossed at least three times to B73. Homozygous wild types (B73) originating from the individual lines, were used as controls. The *bx2* mutants were in a1-m3 background in the W22 inbred line, and the corresponding a1-m3 wild type, also in the W22 background was used as control. The *bx2* mutants and the corresponding wild type (W22) were obtained from Dr. Georg Jander and Kevin Ahern (Boyce Thompson Institute, USA). In homozygous *bx3* plants, no 2,4-dihydroxy-7-methoxy-1,4-benzoxazin-3-one (DIMBOA) can be detected and therefore can be considered null mutants (Frey et al., 1997). Homozygous *bx3* mutants from segregating lines were selected through genotyping using bx3_+540_F (5’-CAC CAA GAA GGT GCA GTC CT-3’), bx3_+1144_R (5’-GTA GCT GGA CTT ACC ACC AAG A-3’), and TIR6 (5’-AGA GAA GCC AAC GCC AWC GCC TCY ATT TCG TC-3’) primers. Genotyping for *tls1* was done as previously described (Durbak et al., 2014). Unless otherwise stated, all maize greenhouse and growth chamber experiments were done with ED73 soil (Einheitserdewerke Werkverband e.V., Sinntal-Altengronau, Germany).

The transgenic Arabidopsis lines, overexpressing parts of the maize benzoxazinoid pathway, namely *pSUR2::Bx1Bx2*, *pSUR2::Bx1Bx2xNahG* and *p35S::Bx3* are described in (Abramov et al., 2021). Seeds of the transgenic lines and the *Col-0* control were sown on 1% H_2_O Agarose plates and stratified for 4 days at 4°C in the dark. Afterwards, seeds were allowed to germinate for 7 days under long day conditions (light: 16 h, 22°C, dark: 8h, 17°C, 40% humidity, light intensity = 65 µE m^-2^ s^-1^), before seedlings were transferred to soil (Floragard B fein, sand, Perligran G mix, 10:1:1).

### Boron concentration measurements

For the GWAS experiment in total 277 inbred lines of the Goodman Buckler association panel (Flint-Garcia et al., 2005) were used. In 2017 and 2018, leaves from the node subtending the ear shoot (referred to as ear leaves) from five plants per inbred line were collected after flowering, pooled, and about 2 cm of the leaves from the tip were discarded. The remaining leaf blades were dried at 60 °C and ground to a fine powder. Boron concentration in individual lines was analyzed subsequently by the azomethine H method (Lohse, 1982) by the Soil and Plant Testing Laboratory at the University of Missouri.

The *bx3* mutants in the B73 background and the B73 inbreds (control) or *bx3* segregating lines were grown in the summers of 2020, 2021, and 2023 at the field station of the University of Bonn (Bonn-Endenich, Germany), where soil boron concentration was 0.27 mg/kg as assessed using a cold 0.01 M CaCl_2_ extraction protocol and a miniaturized curcumin method for analysis (Wimmer and Goldbach, 1998). Four ear leaves of *bx3* mutants, the B73 inbreds or the wild-type sibling controls were pooled separately, 2 cm of the leaf tips were discarded and the remaining leaf blades dried at 60 °C, and ground to a fine powder (three biological replicates). For the Arabidopsis samples, whole rosettes were taken before bolting and dried at 60°C before grinding to a fine powder (three to four biological replicates). Plant samples (500 mg maize or Arabidopsis) were digested with nitric acid in a CEM Mars 5 microwave digestion system (CEM Corporation, Matthews, North Carolina, USA) and boron concentration was analyzed by a miniaturized curcumin method (Wimmer and Goldbach, 1998).

### Genome wide association study (GWAS)

The outlier removal, optimal transformation and Best Linear Unbiased Prediction (BLUP) calculation for the trait were performed as described in (Slaten et al., 2020). The genotypic dataset was previously described in Shrestha et al. (2022). Briefly, the association panel was previously genotyped with the Illumina MaizeSNP50 BeadChip (Cook et al., 2012) and also with a genotyping-by-sequencing approach (Elshire et al., 2011) as described and utilized in Lipka et al. (2013). Single Nucleotide Polymorphisms (SNP)s were filtered using minor allele frequency > 0.05 and a total of 458,775 SNPs from both datasets were used for the GWAS analysis. We used FarmCPU model (Liu et al., 2016) to conduct GWAS and Bonferroni correction was used to correct for multiple testing at 5%. The candidate gene list was obtained using a 200 kb window size (100 kb on either side) of the significant SNPs. This interval was chosen to cover the long range linkage disequilibrium (LD) in maize and also to compensate for low marker coverage as described in (Ching et al., 2002; Flint-Garcia et al., 2003; Yan et al., 2009; Shrestha et al., 2022). The physical locations and annotations of the genes were based on v2 of the maize B73 annotation http://ftp.maizesequence.org/release-5b/filtered-set/.

### Phenotypic analyses

Plant height measurements were taken at maturity from the soil level up to the tip of the tassel (Total plant height = PH) and to the leaf collar of the flag leaf (plant height to flag leaf = FL). In addition, the number of primary tillers was scored. Plant height at seedling stage was determined from the pot soil level to the top of the leaf whorl. Developmental stages were assessed by using V-stages (Abendroth et al., 2011) and leaf number is given as the number of fully developed leaves plus all leaves that have emerged from the whorl.

Tassel length (TL) was determined by measuring the length from the tassel node (node where the tassel is inserted in the stem at the base of the flag leaf) to the tip of the tassel. Peduncle length (PL) was determined by measuring the length of the tassel from the tassel node to the first branch. Branching area (BA) was determined by measuring the length of the main spike between the first and the last branch. Branch number (BN) is the total number of primary tassel branches. The length of the central spike (CS) was determined by subtracting PL and BA from TL.

Boron toxicity symptoms in seedling leaves were assessed using an adapted leaf bronzing score (de Abreu Neto et al., 2017), where 0 = no phenotype, 1 = leaf tips senesced, 2 = leaf tip and edges senesced (with or without leaf rolling), 3 = senescence reaches into leaf blade (with or without leaf rolling), 4 = any or all symptoms of categories 1-3 including aberrant leaf morphology, and 5 = fully senesced leaf (Supplemental Fig. S1).

Percentage of senesced leaf area (boron fertilization experiment) was assessed 14 days after planting (developmental stage V2 with three to four emerged leaves). The entire leaf blade of leaf one was cut and photographed, while for leaf two, 5 cm of the leaf tip were cut and photographed. Image analysis was done with ImageJ (Schneider et al., 2012). The thresholds of all images were adjusted to either provide the full leaf area or the area of senescence (Supplemental Fig. S2). Out of these two values, the percentage of senesced leaf area for every leaf was calculated.

### Boron fertilization

B73 and *bx3* seedlings in the B73 background were grown in ED73 soil in a walk-in growth chamber (26 °C day, 16 °C night, 16 hours day, 8 hours light, 70% humidity) for 14 days and watered with either ultra-pure water (Merck Millipore, Burlington, USA), Peter’s fertilizer (ICL specialty fertilizers, Tel-Aviv, Israel), Peter’s fertilizer + 0.5 mM boric acid, or Peter’s fertilizer + 1 mM boric acid. For the regular strength of Peter’s fertilizer, nitrogen levels were adjusted to 238 ppm, which resulted in a final boron concentration of 0.08 ppm (Matthes et al., 2018). The plants were watered every other day, where 1 l of the respective solution was given to 24 plants (12 plants per genotype). The experiment was repeated three times.

### DIMBOA levels in the *tls1* mutant

The boron transporter mutant *Zmtls1* (Durbak et al., 2014; Leonard et al., 2014) and wild-type siblings were grown in the Sears greenhouse facility at the University of Missouri, Columbia, USA (16/8 h light/dark cycle with an average day temperature of 30.5 °C, an average night temperature of 25 °C, and average humidity of 40% (day) and 60% (night)) and were continuously fertilized with Peter’s fertilizer (ICL specialty fertilizers, Tel-Aviv, Israel), where nitrogen levels were adjusted to 238 ppm, resulting in boron concentrations of 0.08 ppm (Matthes et al., 2018). Developing leaves were dissected as described in (Matthes et al., 2022) and analyzed for 2,4-dihydroxy-7-methoxy-1,4-benzoxazin-3-one (DIMBOA) concentrations using liquid chromatography-mass spectrometry by the proteomics and metabolomics core facility at the University of Nebraska – Lincoln, USA. For details see Supplemental Methods.

### Boron-(H)ION complex formation

A solution of boric acid (2.6 mg/ml in D2O, pH 6, 540 μl, 22.5 μmol, 1.0 eq.) was added to a solution of indolin-2-one (ION, Sigma-Aldrich) (50 mg/ml in DMSO, 60 μl, 22.5 μmol, 1.0 eq.) and kept at room temp. for 80 h. ^11^B-NMR spectra were recorded after mixing, 18 h and 80 h. Further experiments were performed with a boric acid solution, adjusted to pH 8 using 0.5 M NaOH solution, either at room temp. or 80 °C. Spectra were recorded on a Bruker Avance Neo 400 MHz with CryoProbe. Chemical shifts are given in ppm.

A solution of boric acid (2.3 mg/ml in D2O, pH 8 (adjusted using 0.5 M NaOH solution), 540 μl, 20.1 μmol, 1.0 eq.) was added to a solution of 3-hydroxy-ION (HION, Sigma-Aldrich) (50 mg/ml in DMSO, 60 μl, 20.1 μmol, 1.0 eq.) and kept either at room temp. or 80°C. 11B-NMR spectra were recorded after mixing, 3 h, 18 h and 40 h. Further experiments were performed with 10 eq. HION (500 mg/ml in DMSO, 60 μl, 201 μmol, 10 eq.), 0.5 eq. and 0.2 eq. HION (respective stock solutions). The oxidized product isatin was isolated using preparative TLC (CH/AcOEt 1:1).

Control experiments were performed in absence of boric acid and using DMF instead of DMSO, also giving isatin as product.

Under an argon atmosphere, tetrabutylammonium hydroxide (90.0 µL, 184 µmol, 55 w% solution in water, 0.55 eq) was added to a degassed solution of 3-hydroxy-ION (HION, Sigma-Aldrich) (50.0 mg, 335 µmol, 1.00 eq) and boric acid (11.4 mg, 184 µmol, 0.55 eq) in dimethylformamide (15 mL). The resulting solution was stirred at 130 °C for 16 h. Subsequently, the solvent was removed under reduced pressure using a cooling trap (<0.001 mbar, 100°C). The yellow-brown color of the crude product mixture changes immediately to a strong violet color upon contact with air. The ^11^B NMR and ^1^H NMR spectra of the yellow-brown mixture were measured in a quartz glass NMR tube under argon atmosphere, using anhydrous and degassed acetonitrile-*d_3_* as solvent.

To verify the origin of new species in these spectra, the control experiments were performed 1) between the base and boric acid in the absence of HION, 2) between HION and the base in the absence of boric acid, and the crude products were analyzed by ^1^H and ^11^B NMR spectroscopy.

The spectra were recorded on a Bruker Avance 400 NMR spectrometer. Chemical shifts are given in ppm.

### Statistical analysis

Statistical significance of the boron concentrations in the *bx3* mutant, the various growth trait phenotypes between *bx3* and B73 siblings, and the DIMBOA levels in the *tls1* mutant in comparison to non-mutant control lines was assessed in Microsoft Excel using Student’s *t*-test (Student, 1908) at a significance level of *p* < 0.05.

Statistical significance at a significance level of *p* < 0.05 was determined for the different boron fertilization treatments and the different genotypic categories in the Arabidopsis lines using analysis of variance with *post hoc* multiple testing correction applying the Tukey or Benjamini-Hochberg algorithms using the multcompView (Piepho, 2004), agricolae (de Menidburu and Yaseen, 2020), and emmeans (Lenth, 2022) packages in R.

## Results

### Goodman-Buckler association panel shows extensive variation in boron concentration in maize ear leaves

To investigate the extent of phenotypic variability of leaf boron concentrations, boron concentrations from 277 inbred lines (Supplemental Table S1) of the 282 maize Goodman-Buckler association panel (Flint-Garcia et al., 2005) were quantified. The diversity panel was grown in replicates in the summers of 2017 and 2018 at Genetics Farm of the University of Missouri, Columbia, USA, where soil boron concentrations of 0.39 mg/kg are considered low (Durbak et al., 2014; Matthes et al., 2018). This can be seen by the occurrence of the strong boron deficient leaf phenotype of the maize *Zmtassel-less1* (*Zmtls1*) mutant, which is defective in active boron transport (Durbak et al., 2014; Matthes et al., 2022).

After flowering, the boron concentration of ear leaves (the leaf subtending the ear shoot) was analyzed. In the analyzed back-transformed best linear unbiased predictors (BLUPs) from the diversity panel, the boron concentration varied between 8.6 µg/g (line CMV3) and 17.8 µg/g (line IL14H) (Supplemental Table S1) with a mean of 11.9 µg/g (Supplemental Table S2). The broad sense heritability calculated on a line-mean basis was 0.31 (Supplemental Table S2), indicating that the boron concentration in maize ear leaves is a complex trait and warrants genetic dissection to unravel this complexity.

### GWAS uncovers two significant loci associated with the natural variation in boron concentration

To associate the observed variation in ear leaf boron concentration with the diverse inbred lines in the Goodman-Buckler association panel, we performed a genome wide association study (GWAS) using BLUPs from the analyzed 277 inbred lines. Using the publicly available SNP data from the 282 Goodman-Buckler association, two significant SNP-trait associations (Fig. 1A) were identified. One SNP was on chromosome four (chr4) and one on chr7 (Fig. 1A, Table 1), suggesting that the genomic regions that are in linkage disequilibrium (LD) with them were significantly associated with the ear leaf boron concentration in maize. A quantile-quantile plot from the GWAS showed our GWAS model is not inflated and depicts that few *p*-values of the performed association tests between SNP data and boron concentration data were more significant than expected (Fig. 1B).

**Figure 1:**
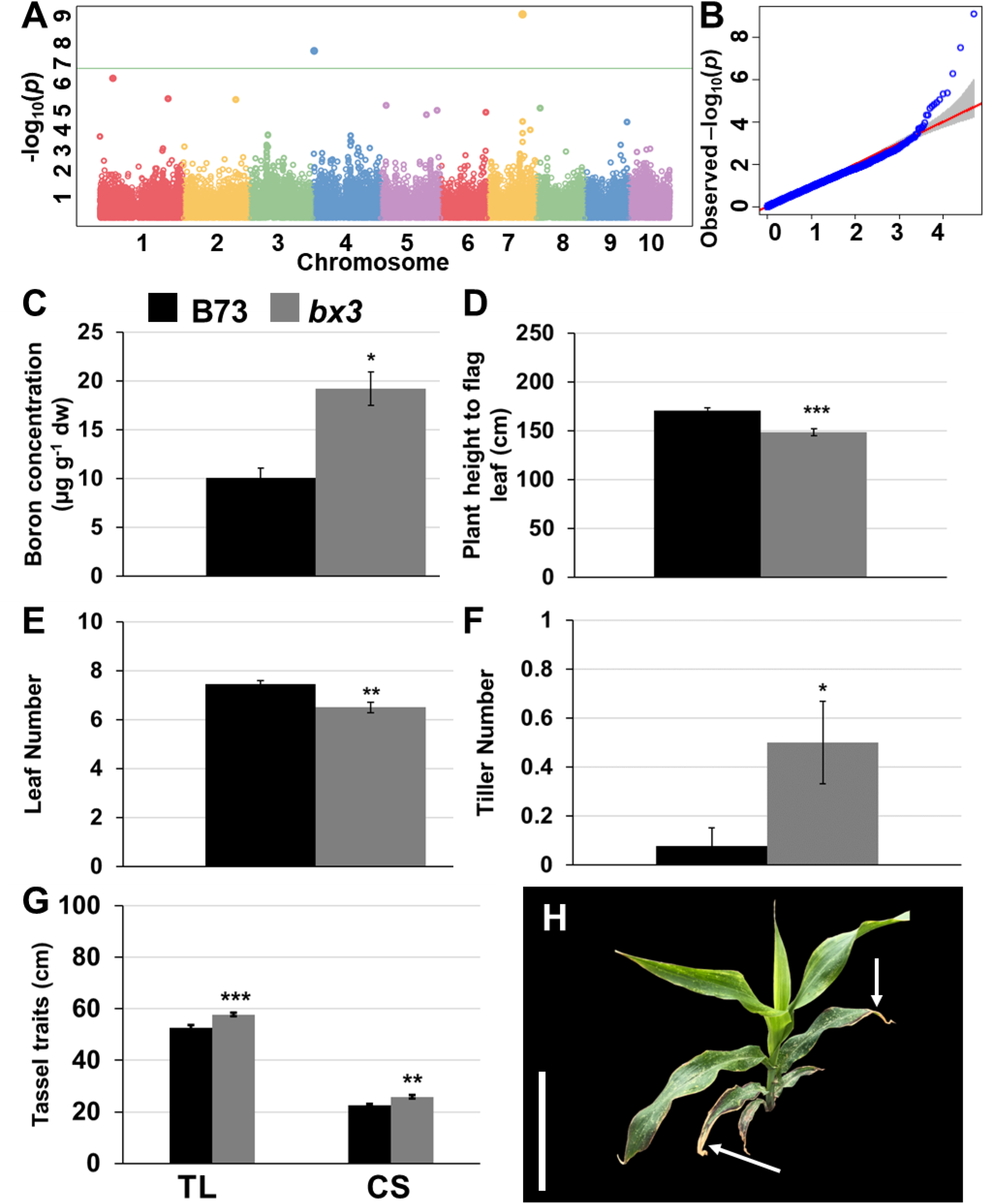
Phenotypes of the *bx3* mutant grown in the field (Bonn-Endenich, 2020). A) Manhattan plot depicting genomic regions associated with variations of boron concentrations in a subset of the 282 Goodman-Buckler association panel. B) Quantile-quantile (qq) plot depicting the expected (x-axis) and observed (y-axis) – log(10) *p*-values. C) Boron concentration in ear leaves of the *bx3* mutant and B73 siblings. D) Plant height to flag leaf at maturity of the *bx3* mutant and B73 siblings. E) Leaf number between the leaf subtending the ear and the flag leaf of the *bx3* mutant and B73 siblings. F) Tiller number of the *bx3* mutant and B73 siblings. G) Tassel traits of the *bx3* mutant and B73 siblings. H) *bx3* plant image at 42 days after planting. Note arrows in H) depicting leaf senescence at the leaf blade tip and the margins of *bx3*. Statistical analyses in C through G depict averages with standard error of means. Sample numbers in C – F were 13 (B73) and 14 (*bx3*) and in G 10 (B73) and 11 (*bx3*). Statistical significance was calculated using Student’s *t*-Test (* *p* < 0.05, *** *p* < 0.005). Scale bar in H = 15 cm. TL = tassel length, CS = length of central spike.

**Table 1:**
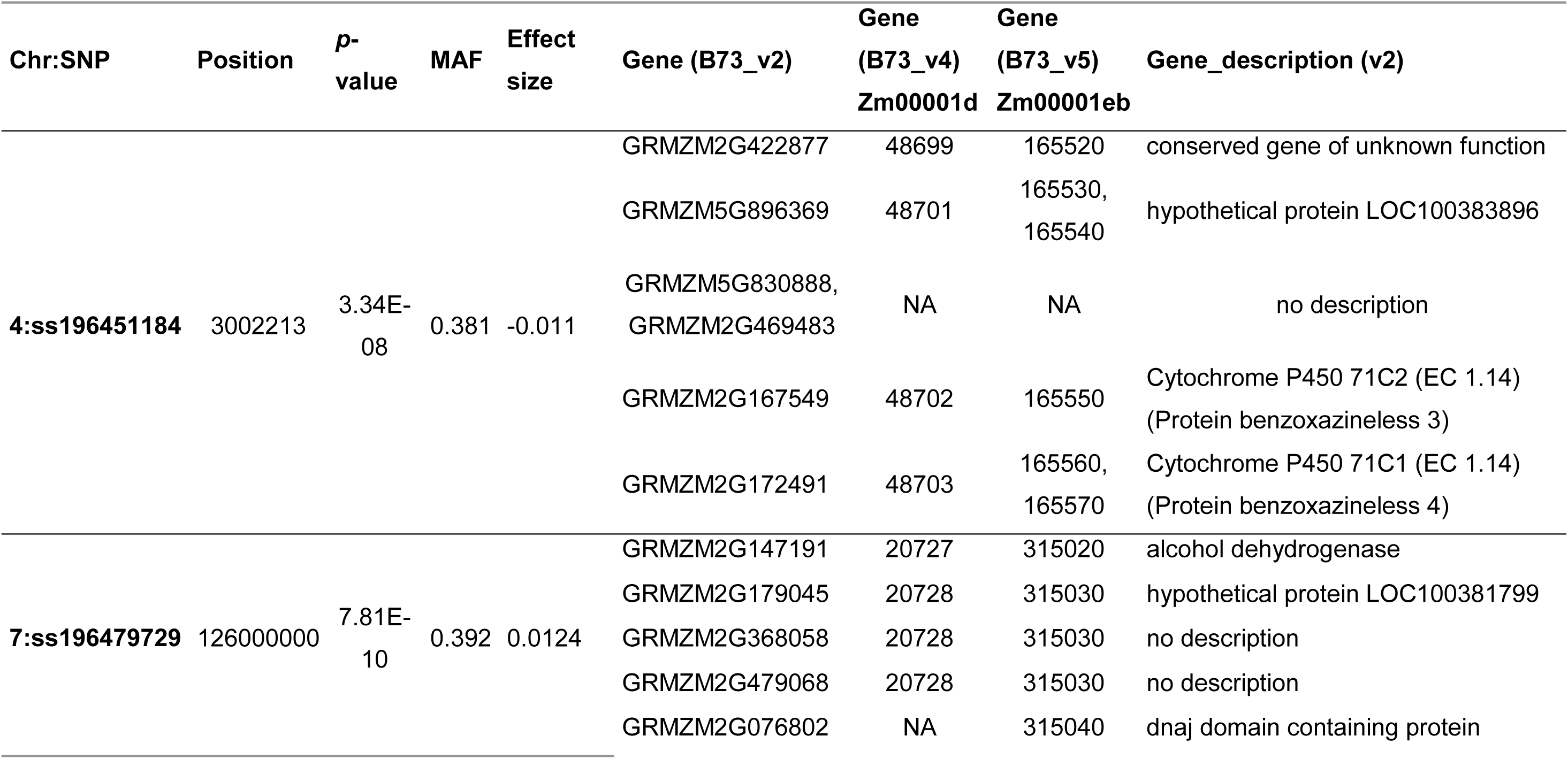
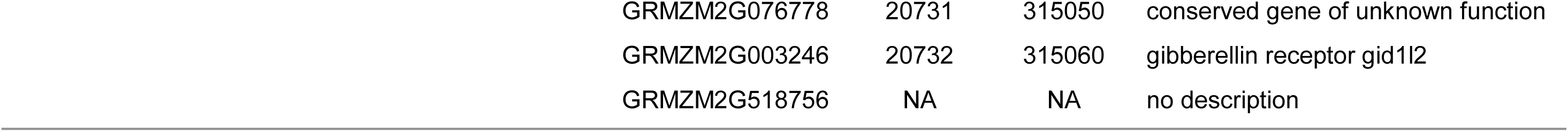
GWAS of the boron concentration in ear leaves of the 282 Goodman-Buckler association panel and the candidate genes from 200 kb interval centered on the significant SNP (100 kb upstream/ downstream) are summarized. Chromosome (Chr.), SNP ID, position, *p*-value, minor allele frequency (MAF), effect size, gene ID based on V2, V4, and V5 assembly and gene description are summarized.

| Chr:SNP | Position | *p*-value | MAF | Effect size | Gene (B73_v2) | Gene (B73_v4)<br>Zm00001d | Gene (B73_v5)<br>Zm00001eb | Gene_description (v2) |
| --- | --- | --- | --- | --- | --- | --- | --- | --- |
| 4:ss196451184 | 3002213 | 3.34E-08 | 0.381 | -0.011 | GRMZM2G422877 | 48699 | 165520 | conserved gene of unknown function |
|  |  |  |  |  | GRMZM5G896369 | 48701 | 165530,<br>165540 | hypothetical protein LOC100383896 |
|  |  |  |  |  | GRMZM5G830888,<br>GRMZM2G469483 | NA | NA | no description |
|  |  |  |  |  | GRMZM2G167549 | 48702 | 165550 | Cytochrome P450 71C2 (EC 1.14)<br>(Protein benzoxazineless 3) |
|  |  |  |  |  | GRMZM2G172491 | 48703 | 165560,<br>165570 | Cytochrome P450 71C1 (EC 1.14)<br>(Protein benzoxazineless 4) |
| 7:ss196479729 | 126000000 | 7.81E-10 | 0.392 | 0.0124 | GRMZM2G147191 | 20727 | 315020 | alcohol dehydrogenase |
|  |  |  |  |  | GRMZM2G179045 | 20728 | 315030 | hypothetical protein LOC100381799 |
|  |  |  |  |  | GRMZM2G368058 | 20728 | 315030 | no description |
|  |  |  |  |  | GRMZM2G479068 | 20728 | 315030 | no description |
|  |  |  |  |  | GRMZM2G076802 | NA | 315040 | dnaj domain containing protein |

|  |  |  |  |
| --- | --- | --- | --- |
| GRMZM2G076778 | 20731 | 315050 | conserved gene of unknown function |
| GRMZM2G003246 | 20732 | 315060 | gibberellin receptor gid1l2 |
| GRMZM2G518756 | NA | NA | no description |

We identified six candidate genes from the 200 kb intervals (100 kb up and down stream) centered around the significant SNP on chr4 and eight candidate genes centered around the SNP on chr7 (Table 1). Two of the six chr4 genes were annotated as *benzoxazinless3* (*bx3*) and *bx4* (Table 1), involved in benzoxazinoid biosynthesis. The annotated genes underlying the peak on chr7 encoded an alcohol dehydrogenase, a DNAJ domain containing protein, and the gibberellin receptor GID1L2 (Table 1).

### Correlation analysis of gene expression of GWAS hits and boron concentration in the Goodman-Buckler association panel

To assess a potential connection between the identified GWAS candidate genes’ transcript expression and boron concentration in the maize ear leaf, a correlation analysis was performed using publicly available gene expression data of the GWAS candidate genes in leaf tissues of the 282 Goodman-Buckler association panel (Kremling dataset, Kremling et al., 2018) (Supplemental Table S3; Supplemental Methods). Of the 14 detected GWAS hits, data for six genes were available from the Kremling dataset. We detected significant correlations between gene expression and boron concentration for five of the GWAS hits (*bx3* on chr4; alcohol dehydrogenase, DNAJ domain containing protein, GRMZM2G076778 (which is a gene with unknown function), and Gibberellin Receptor on chr7) in one or multiple leaf tissues, suggesting that the variation of boron concentration in the Goodman-Buckler association panel might be correlated with expression level changes of these candidate genes. While *bx3* and the alcohol dehydrogenase gene showed a negative correlation in the third leaf, the DNAJ domain containing protein and the Gibberellin Receptor genes showed positive correlations in various leaf tissues. GRMZM2G076778 depicted a negative correlation in adult leaf tissue (Supplemental Table S3).

### The GWAS candidate genes *bx3* and *bx4* are differentially expressed in the maize boron transporter mutant *Zmtls1*

To further assess a connection between the identified genes and boron concentration in maize leaves, we tested, whether the identified GWAS candidate genes are differentially expressed in the maize boron transporter mutant *Zmtls1*, for which RNA-seq data of developing tassel meristems (∼ 1 mm) are publicly available (Matthes et al., 2022). In *Zmtls1* tassel meristems, boron levels were found to be reduced compared to wild-type siblings (Durbak et al., 2014). We found that the chr4 candidate genes *bx3* and *bx4* were significantly upregulated in *Zmtls1* in that dataset, with log_2_ fold changes of 1.56 for *bx3* (*p-adj* = 0.0013) and 1.42 for *bx4* (*p-adj* =0.0025) (Matthes et al., 2022) (Table 2). Notably, none of the other candidate genes on chr4 or chr7 appeared as differentially expressed in the *Zmtls1* RNA-seq data set (Table 2).

**Table 2:** Differential expression of GWAS candidate genes and benzoxazinoid biosynthesis genes in developing meristems of *Zmtls1* compared to wild-type controls. ns = not statistically significantly different.

|  | *tls1* vs<br>WT<br>(log<sub>2</sub>FC) | *p*-adj |
| --- | --- | --- |
| Zm00001d048702 (*bx3*) | 1.56 | 0.001288 |
| Zm00001d048703 (*bx4*) | 1.42 | 0.002502 |
| Zm00001d48699 (unknown) |  | ns |
| Zm00001d48701 (hypothetical) |  | ns |
| Zm00001d20727 (alcohol dehydrogenase) |  | ns |
| Zm00001d20728 (hypothetical) |  | ns |
| Zm00001d20731 (unknown) |  | ns |
| Zm00001d20732 (gibberellin receptor<br>gid1l2) |  | ns |
| Zm00001d048709 (*bx1*) |  | ns |
| Zm00001d048710 (*bx2*) |  | ns |
| Zm00001d048705 (*bx5*) |  | ns |
| Zm00001d048634 (*bx6*) | 1.66 | 1.66E-16 |
| Zm00001d049179 (*bx7*) |  | ns |
| Zm00001d048707 (*bx8*) |  | ns |

Taken together, the chr4 candidate gene *bx3* was the only GWAS candidate, whose expression in leaf tissue (Kremling et al., 2018) showed a significant correlation with the measured leaf boron concentration in the 282 Goodman-Buckler association panel (negative correlation) and which was also differentially expressed in the boron-deficient tassel meristem of the *Zmtls1* mutant.

### Mature plants of maize *bx3* mutants have a boron concentration phenotype

The previous GWAS and expression analyses linked the *bx3* gene with variation in boron concentration in maize. To test if there was a boron associated phenotype in *bx3* mutants, we analyzed boron concentration in ear leaves of mature *bx3* mutants (Frey et al., 1997) and in the B73 inbred line grown at the field site in Bonn, where soil boron levels averaged 0.27 mg/kg. Soil boron concentrations in combination with the appearance of the strong seedling phenotype of the boron deficient maize mutant *Zmtls1* in Bonn-Endenich (Supplemental Fig. S3) showed that these field site conditions are comparable to those at the Genetics Farm in Columbia, Missouri (Durbak et al., 2014), the field site used for the GWAS.

Analyses from field grown plants in 2020 and 2021 showed that boron concentration was significantly elevated in the ear leaves of *bx3* mutants compared to B73 (Fig. 1C, Supplemental Tables S4, S5), substantiating a correlation between *bx3* and boron levels in maize. This increase was nutrient specific as the concentrations of the other tested micro- and macronutrients were either unchanged (magnesium, calcium, phosphorus, sulfur, iron, zinc) or even decreased (manganese, copper) in the *bx3* mutant. Only potassium also showed a mild increase in *bx3* ear leaves (Supplemental Table S4).

During the 2023 field season, however, no elevated boron concentrations were detected in *bx3* ear leaves compared to B73 (Supplemental Fig. S3). Similar observations were made in the greenhouse, when *bx3* mutants were grown in boron-rich soil (ED73 soil boron = 1.97 mg/kg) (Supplemental Fig. S4). In addition, the *Zmtls1* phenotype was also less severe with enhanced boron concentrations in ear leaves under the 2023 field conditions and under greenhouse conditions (Supplemental Fig. S3). It, therefore, seems likely that either variation in soil boron concentration or additional, not yet identified environmental factors influenced the boron-related phenotype in *bx3* and *tls1*.

Elevated boron concentrations in *bx3* ear leaves compared to B73 siblings did not lead to striking differences in reported boron toxicity phenotypes (Supplemental Fig. S4A,B), like leaf tip necrosis in mature leaves (Supplemental Fig. S5), suggesting that the elevated boron concentrations in *bx3* ear leaves were not toxic. However, earlier during vegetative development (42 days after planting in the field), *bx3* mutants showed severe leaf senescence at the leaf blade tip of particularly older leaves, extending into the leaf blade margins, which was not observed in B73 (arrowed in Fig. 1H, Supplemental Fig. S5B), suggesting a developmental factor influencing the phenotypes observed in *bx3* mutants.

In addition, the *bx3* mutants showed minor alterations of other potentially boron related phenotypes: Plant height (up to the flag leaf) was significantly reduced in *bx3* mutants compared to B73 (Fig. 1D), and *bx3* mutants had significantly fewer leaves between the ear and the tassel compared to B73 siblings (Fig. 1E). In addition, tiller number, tassel length, and the length of the central spike were significantly increased in *bx3* mutants compared to B73 siblings (Fig. 1F – G, Supplemental Table S5). While kernel row number was slightly, but significantly reduced in *bx3* mutants compared to B73 control plants, ear length was significantly longer (Supplemental Table S5). These morphological data suggested that enhanced boron levels in the *bx3* mutants did not lead to striking boron toxicity symptoms in field grown plants at maturity.

### *bx3* mutants show a leaf tip necrosis phenotype during seedling development and its severity correlates with boron levels

In order to characterize the impact of boron levels on the observed leaf tip necrosis phenotype that we observed six weeks after planting (Fig. 1H), we analyzed the phenotype of *bx3* mutants earlier in development grown with different levels of boron. We first grew *bx3* mutants and B73 siblings in the growth chamber in boron sufficient soil (boron concentration of ED73 soil = 1.97 mg/kg) and phenotypically characterized seedling development over time (Fig. 2, Supplemental Table S6). We evaluated plant height, V-stage, leaf tip necrosis and leaf number at 14 days (developmental stage: V1/2 with three to four leaves emerged) and 25 days after planting (DAP) (developmental stage: V3/4, with five to six leaves emerged) (Fig. 2; Supplemental Fig. 1G, Supplemental Table S6). At both time points, the *bx3* mutant was phenotypically not distinguishable from B73 with respect to plant height, leaf number, and V-Stage (Supplemental Table S6). However, at 14 DAP (developmental stage: V1/2), when both B73 and *bx3* had three to four leaves, the leaf tips of leaf 1 and 2 showed a distinct leaf tip necrosis phenotype in *bx3* mutants, but not in the B73 control (Fig. 2A-C). In addition, we detected statistically significantly higher boron concentrations in pooled first and second leaves of *bx3* mutants compared to those of B73 14 DAP (Fig. 2D). This showed, that boron levels are also enhanced in seedling leaves of the *bx3* mutant and suggested that these elevated boron levels correlated with the observed leaf tip necrosis phenotype.

**Figure 2:**
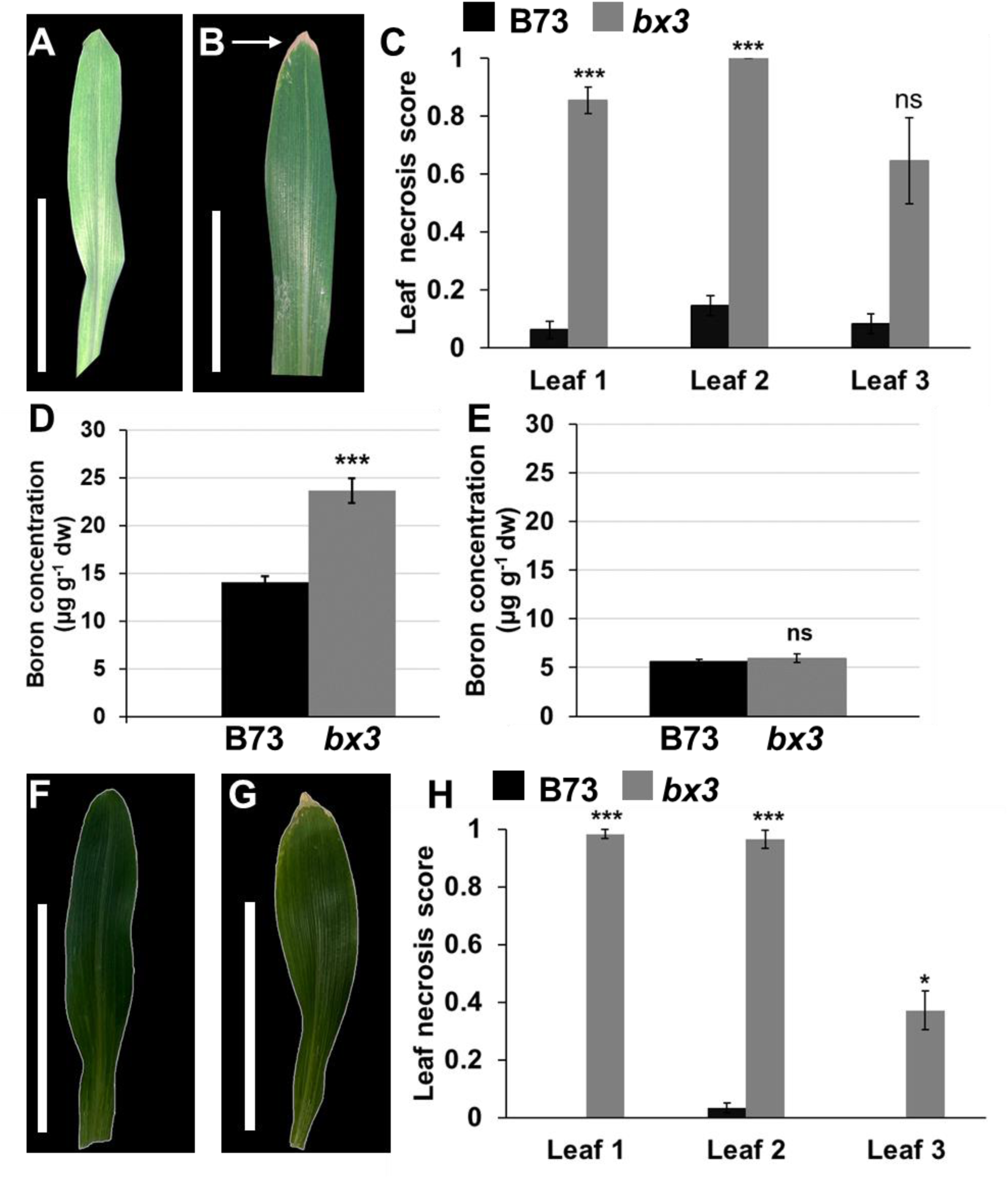
Leaf phenotypes of B73 and *bx3* seedlings. Images of the first leaves of A) B73 and B) *bx3* 14 days after planting (DAP) grown in the growth chamber in ED73 soil (boron concentration = 1.97 mg/kg). Arrows indicate leaf tip senescence. C) Statistical analysis of leaf tip necrosis in all emerged leaves of ED73-grown B73 and *bx3* 14 DAP. Depicted are means of three biological replicates, where n = 16 per genotype and biological replicate. Error bars show standard error of means. Statistical analysis of boron concentration (µg g^-1^ dw) in leaf one and two (pooled) of B73 and *bx3* 14 DAP, when D) seedlings grew in ED73 soil and E) seedlings grew in HGoTech substrate (boron concentration 0.03 mg kg^-1^). Given are means over four biological replicates. The error bars represent standard error of means. Statistical significance at * *p* < 0.05, ** *p* < 0.01, *** *p* < 0.005 according to Student’s *t*-test. Images of the first leaves of F) B73 and G) *bx3* 14 DAP grown in the growth chamber in HGoTech substrate. H) Statistical analysis of leaf tip necrosis in all emerged leaves of HGoTech-grown B73 and *bx3* 14 DAP. Depicted are means of four biological replicates, where n = 15 per genotype and biological replicate. Error bars show standard error of means. Scale bars in A, B, F, G = 5 cm. ns = statistically not significantly different.

Since the *Zmtls1* mutant showed enhanced boron levels in ear leaves under field conditions in 2023 (Supplemental Fig. S3), we did not use a genetics approach using double mutant analysis to determine the effects of lowered boron levels on *bx3* mutants. Instead, we grew the *bx3* mutants and B73 control plants in boron free media (HGoTech GmbH, Bonn, Germany), where boron concentration = 0.03 mg/kg.

The leaf tip necrosis phenotype remained present in the *bx3* mutant, when grown in the boron-free media, whereas boron concentration in seedling leaves of *bx3* (leaves 1 and 2 pooled) were not statistically significantly different from the B73 control (Fig. 2 E-H). Therefore, we tested whether adding extra boron could influence the severity of the leaf tip necrosis phenotype by subjecting *bx3* mutants and B73 control plants (grown in ED73 soil) to the following watering regimes: Ultra-pure water (Merck Millipore, Burlington, USA), Peter’s fertilizer (ICL specialty fertilizers, Tel Aviv, Israel), Peter’s fertilizer + 0.5 mM boric acid, and Peter’s fertilizer +1 mM boric acid. The ultra-pure water and the Peter’s fertilizer treatments showed the leaf tip necrosis phenotype in *bx3* plants, but not in B73 plants 14 DAP. In contrast, the treatments with Peter’s fertilizer and additional boric acid caused the occurrence of the leaf tip necrosis phenotype also in B73 seedlings (Fig. 3A-C, Supplemental Table S7), suggesting that the additional boric acid treatment caused this phenotype even in B73. In the *bx3* mutant, no enhancement of the leaf tip necrosis score was detected between the different watering regimes, however the senesced leaf area in *bx3* mutants was significantly larger in the additional boric acid treatments compared to the ultra-pure water and Peter’s fertilizer treatments (Fig. 3D).

**Figure 3:**
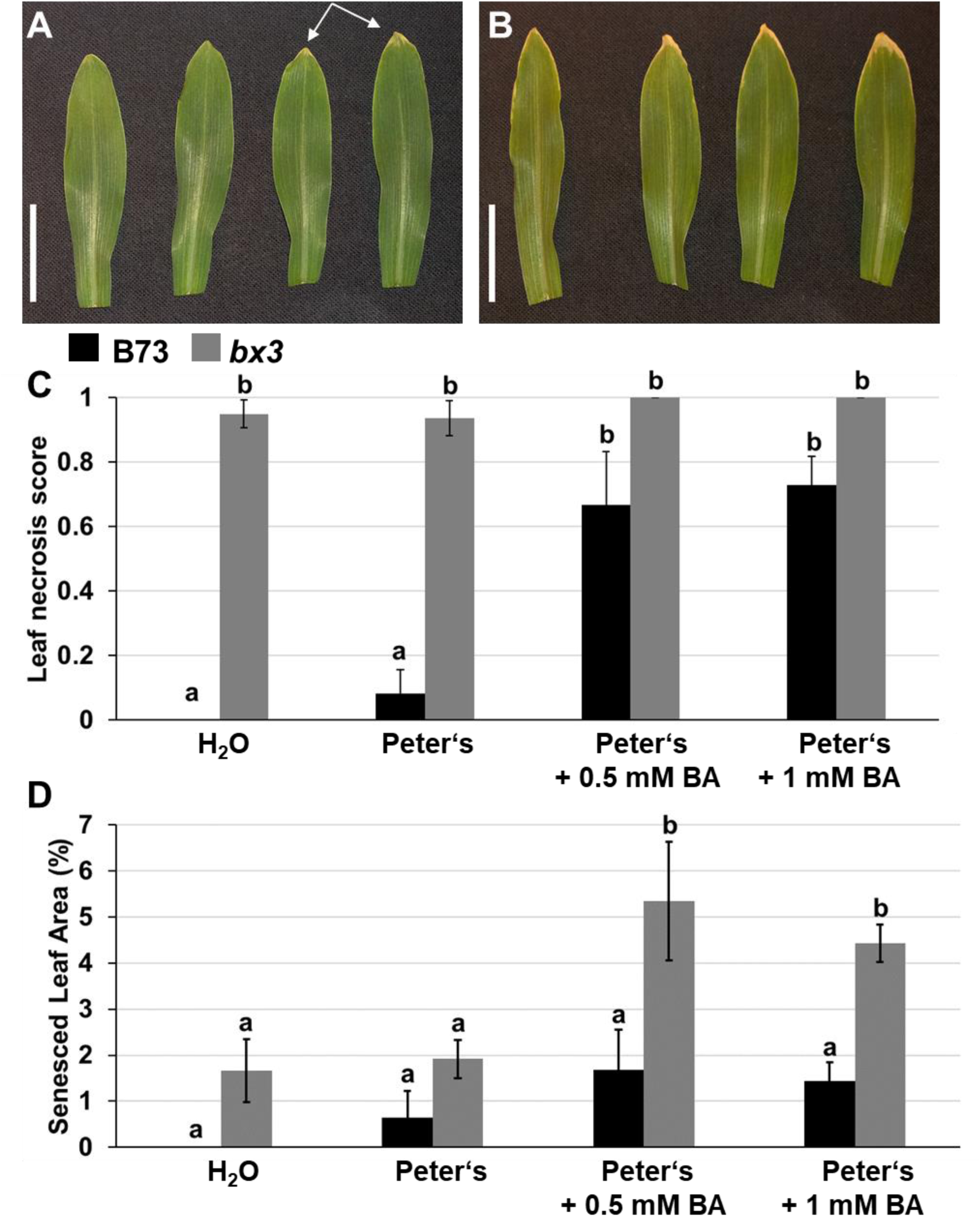
Senesced leaf area is enhanced by boron fertilization. Images of first leaves of A) B73 and B) *bx3* grown in ED73 soil. The images were taken 14 days after planting (DAP) with different watering regimes (from left to right: H_2_O, Peter’s fertilizer, Peter’s fertilizer + 0.5 mM boric acid (BA), Peter’s fertilizer + 1 mM BA). Arrows in A point to leaf tip necrosis in B73. C) Statistical analysis of leaf necrosis score 14 DAP in B73 and *bx3* seedlings. D) Statistical analysis of senesced leaf area 14 DAP in B73 and *bx3* seedlings. C and D show means of four biological replicates with 4-6 individuals per genotype and replicate. Error bars represent standard error of means. Different letters indicate statistical significance at *p* < 0.05 according to an analysis of variance with *post hoc* multiple comparison correction using the Tukey algorithm.

Taken together, these analyses showed that the severity of the leaf tip necrosis phenotype in *bx3* mutants is correlated with enhanced boron levels. However, the variability of the results indicated there are also additional, so far unidentified factors, contributing to the leaf tip necrosis phenotype in *bx3* mutants.

### Investigation of a correlation between additional benzoxazinoid pathway genes and boron homeostasis

To determine if the link between boron levels and *bx3* was specific to *bx3*, we investigated other genes in the benzoxazinoid pathway. *bx3* encodes a cytochrome P450 monooxygenase that is involved in the biosynthesis of benzoxazinoids, a group of specialized secondary metabolites (Frey et al., 1997) (Supplemental Fig. S6). The predominant benzoxazinoids are 2,4-dihydroxy-1,4-benzoxazin-3-one (DIBOA) and its C-7-methoxy derivative DIMBOA, the latter being the major benzoxazinoid in B73 maize. There are 14 characterized *bx* genes in maize (as reviewed in de Bruijn et al., 2018) with *bx1* to *bx9* involved in the synthesis of DIMBOA and *bx1* to *bx8* being located in proximity on chr4 (as reviewed in Frey et al., 2009). We therefore assessed the possibility that the GWAS detected a significant correlation with the *bx* gene cluster on chr4, rather than individual *bx* genes, which was also reasoned by the detection of both *bx3* and *bx4* in the GWAS (Table 1). We performed correlation analyses with publicly available expression data in leaf tissue (Kremling et al., 2018) for all *bx* genes located on chr4 and the boron concentration data obtained from 277 lines of the 282 Goodman-Buckler association panel (Supplemental Table S1), similar to the correlation analyses with the detected GWAS candidates (Supplemental Table S3). These analyses showed that, in addition to gene expression of *bx3*, that of *bx2* also showed a negative correlation in the base of third leaf tissue, and that of *bx5* showed a positive correlation in the tip of third leaf tissue with boron concentration (Supplemental Table S8).

Furthermore, we examined *bx* gene expression of the chr4 cluster in the developing tassel meristem data set of the *Zmtls1* mutant, which has reduced boron levels in that tissue. We found that in addition to *bx3* and *bx4*, also expression of *bx6* is significantly upregulated in *Zmtls1* mutants compared to normal siblings (Table 2). These results indicated that a potential correlation between *bx* genes and boron concentration in maize might not be restricted to *bx3* and that benzoxazinoid biosynthesis could be altered in *Zmtls1*. Therefore, we quantified levels of DIMBOA in *Zmtls1* leaves and found that the levels were significantly reduced compared to wild type controls (WT = 462.01 ± 71.01 ng/g, *Zmtls1* = 229.41 ± 59.99 ng/g, *p* < 0.05).

### Boron homeostasis appears to be linked to *bx3* rather than DIMBOA

To test potential correlations between boron levels and benzoxazinoid biosynthesis, we analyzed boron concentration and assessed the leaf tip necrosis phenotype in *bx1* and *bx2* mutants, which were reported to have reduced DIMBOA levels (Hamilton, 1964; Frey et al., 1997; Tzin et al., 2015).

We found, that boron levels in ear leaves of *bx1* or *bx2* mutants were not statistically significantly different compared to their respective inbred control (Fig. 4A), when grown in Bonn-Endenich (soil boron concentration: 0.27 mg/kg in 2021). Likewise, *bx1* (developmental stage: V1/2) and *bx2* mutants (VE/1) did not show the striking leaf tip necrosis phenotype observed in *bx3* mutants (developmental stage: V1/2) at 14 DAP (Fig. 4B - E, Supplemental Table S9). While the *bx2* mutant (introgressed in W22 and not B73) was not at the same developmental stage (VE/1) as *bx1* or *bx3* at 14 DAP, it did not show a statistically significant difference in leaf tip necrosis compared to its inbred control (W22) in any of the analyzed timepoints (Supplemental Table S9). In contrast, *bx1* showed a similar leaf tip necrosis phenotype to *bx3* at 18 DAP (developmental stage of both mutants: V2) and at 25 DAP (developmental stage of both mutants: V3). This phenotype, however was restricted to the first leaf (18 DAP) or the first two leaves (25 DAP) (Fig. 4, Supplemental Table S9), which was in contrast to *bx3*, where the leaf necrosis phenotype was present in all developed leaves (Supplemental Table S9).

**Figure 4:**
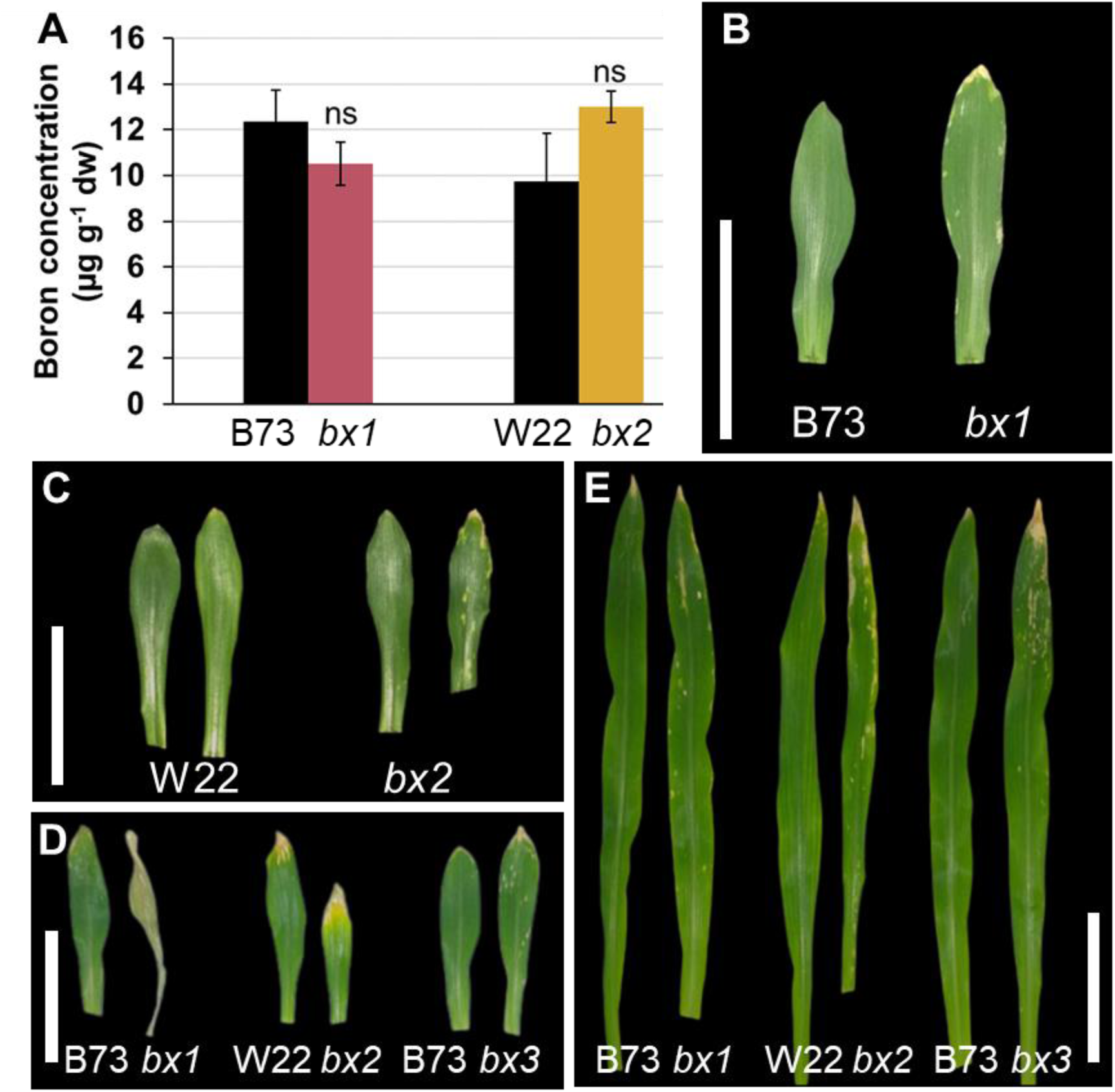
Boron related phenotypes observed in the *bx3* mutant are not found in *bx1* or *bx2*. A) Statistical analysis of boron concentration (µg g^-1^) in ear leaves of *bx1* and *bx2* mutants compared to the respective inbred control (grown in the field, Bonn-Endenich). Depicted are means ± standard error of means and statistical significance was calculated using Student‘s *t-test* (*p* < 0.05) (n = 3, where each replicate is a pool of 4 individual leaves.) Leaf phenotypes of B73, W22, and mutants of *bx1*, *bx2*, *bx3* at 14, 18, and 25 days after planting (DAP), where B) Leaf 1 at 18 DAP, C) Leaf 1 at 18 DAP, D) Leaf 1 at 25 DAP, E) Leaf 2 at 25 DAP. Scale bars in B - E = 5 cm.

Taken together, our analyses therefore suggested a link between boron and *bx3* rather than boron and DIMBOA and raised the question whether the lack of a functional BX3 enzyme or the accumulation of the BX3 substrate (Supplemental Fig. S6A) is connected with the observed boron accumulation.

### Boron concentration is elevated in transgenic Arabidopsis lines overexpressing BX1 and BX2

Although all three maize *bx* mutants we investigated are DIMBOA deficient (Frey et al., 1997; Tzin et al., 2017), *bx3* unlike *bx1* and likely *bx2* also accumulates the intermediate indolin-2-one (ION), which is the substrate of the BX3 enzyme (Abramov et al., 2021; Supplemental Fig. S6A). In order to test, whether elevated ION levels correlate with the observed boron concentration elevation in the *bx3* mutant, we made use of Arabidopsis lines, where parts of the benzoxazinoid biosynthesis pathway were transgenically introduced (Abramov et al., 2021). Arabidopsis does not endogenously express this pathway, allowing an assessment of a potential boron-benzoxazinoid correlation in a ‘clean’ background. We found, that boron concentration in rosette leaves at bolting stage of *pSUR2::Bx1Bx2*, which accumulate ION, was significantly higher compared to the *Col-0* control. On the other hand, functional expression of BX3 (*p35S*::Bx3 lines, which do not accumulate ION) did not influence the boron concentration. Therefore, a direct effect of the enzyme seems to be unlikely (Fig. 5). In addition, *pSUR2::Bx1Bx2* overexpressing lines showed a subtle leaf tip necrosis phenotype, compared to *Col-0* and *p35S::Bx3* lines (Fig. 5). Since *pSUR2::BX1BX2* lines were reported to additionally show elevated salicylic acid (SA) levels, we additionally assessed the impact of SA accumulation on the boron related phenotypes of *pSUR2::Bx1Bx2* lines, by analyzing *pSUR2::Bx1Bx2* crossed to *NahG* lines (*pSUR2::Bx1Bx2 x NahG*), where the SA hydroxlase *NahG* was introduced (Abramov et al., 2021). Our results showed, that boron concentration is also significantly elevated in *pSUR2::Bx1Bx2xNahG* rosette leaves compared to *Col-0* controls and that the leaf tip necrosis phenotype in these lines is enhanced (Fig. 5E – F, Supplemental Table S10). Therefore, these results suggested, that a correlation between boron levels and the benzoxazinoid pathway is linked to an accumulation of the intermediate ION.

**Figure 5:**
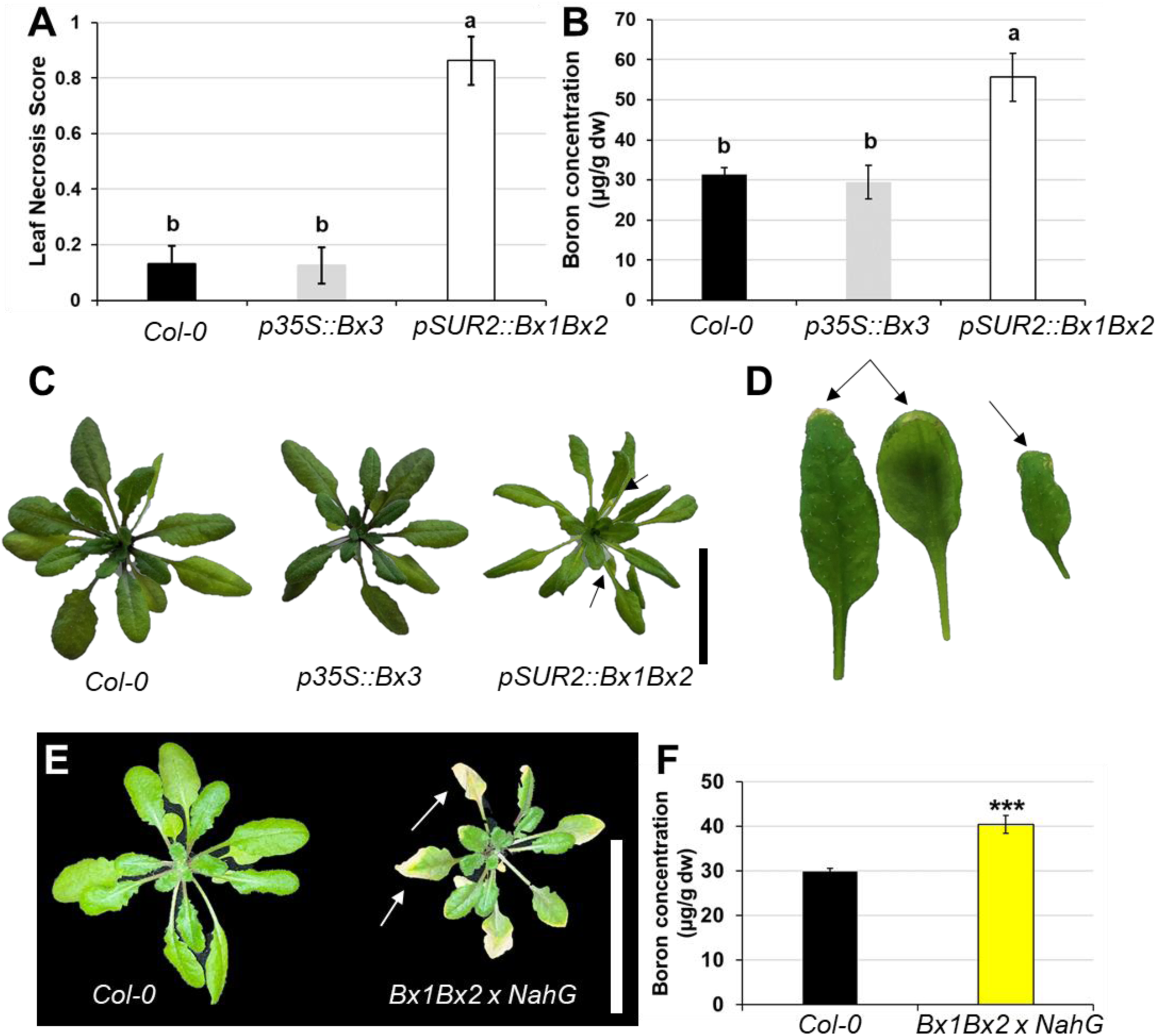
Analysis of boron-benzoxazinoid relations in transgenic Arabidopsis. A) Statistical analysis of leaf tip necrosis in *Col-0*, *p35S::Bx3* and *pSUR2::Bx1Bx2* Arabidopsis lines. B) Statistical analysis of boron concentration (µg g^-1^ dw) in *Col-0*, *p35S::Bx3*, and *pSUR2::Bx1Bx2* Arabidopsis lines. C) Rosette leaf phenotypes of *Col-0*, *p35S::Bx3*, *pSUR2::Bx1Bx2*. Note arrows in *pSUR2::Bx1Bx2* pointing to a leaf tip necrosis phenotype in older leaves. D) Images of different leaves showing a subtle leaf tip necrosis phenotype in *pSUR2::Bx1Bx2* transgenic Arabidopsis. Note arrows that point to the leaf tip necrosis phenotype. E) Rosette leaf phenotypes of *Col-0* and *pSUR2::Bx1Bx2* in the *NahG* mutant background (*pSUR2::Bx1Bx2xNahG*). Note arrows in *pSUR2::Bx1Bx2xNahG* pointing to severely senesced rosette leaves. F) Statistical analysis of boron concentration (µg g^-1^ dw) in *Col-0* and *pSUR2::Bx1Bx2xNahG* Arabidopsis lines. Statistical analyses in A, B, and F depict means over 3-4 biological replicates (n = 13-15 individuals per biological replicate) ± standard error of means. Different letters indicate statistically significant differences according to an analysis of variance and *post hoc* Tukey test (*p* < 0.05). *** indicates statistical significance at *p* < 0.005 according to Student‘s *t*-test. Scale bars in C = 4.5 cm, in E = 5 cm.

### Boric acid forms a complex with the benzoxazinoid intermediate 3-hydroxy-ION (HION)

The benzoxazinoid DIMBOA was reported to chelate iron thus making it bioavailable (Hu et al., 2018), but also complex formation between DIMBOA and other nutrients, including zinc, copper and manganese was reported (as reviewed in Wouters et al., 2016). We therefore reasoned that elevated boron levels in the *bx3* mutant could be related to a benzoxazinoid-mediated alteration of boron transport or mobility. The correlation between boron levels and an accumulation of ION led us to test, the hypothesis that boric acid and ION might form a complex, which in turn might influence boron transport or mobility in maize. The reaction between ION and boric acid however, did not lead to any boric acid-ION complex (Supplemental Fig. S7). As boron is known to specifically interact with *cis*-diol groups, which are not present in the benzoxazinoid intermediate ION, we also tested the next intermediate in the benzoxazinoid pathway, 3-hydroxy-ION (HION). HION is the product of the enzymatic function of BX3 (Supplemental Fig. S6A) that may tautomerize to the lactim species bearing two *cis*-diol groups (Supplemental Fig. S6B). The reaction of HION with boric acid in the presence of a base led to numerous additional signals in the ^11^B NMR spectrum (Fig. 6A) and ^1^H NMR spectrum (Supplemental Fig. S8A). Importantly, these signals were not observed when the analysis was done with boric acid and base alone (Fig. 6B) or when the control experiment was performed in the absence of boric acid (Supplemental Fig. S8B). These results suggested the formation and therefore the existence of additional boron compounds formed between boric acid and HION (Fig. 6). Attempts to identify the nature of these additional boron compounds were unsuccessful, suggesting that such a complex is unstable. In the presence of air, HION reacted to isatin, which was not dependent on the presence of boric acid (Supplemental Fig. S7).

**Figure 6:**
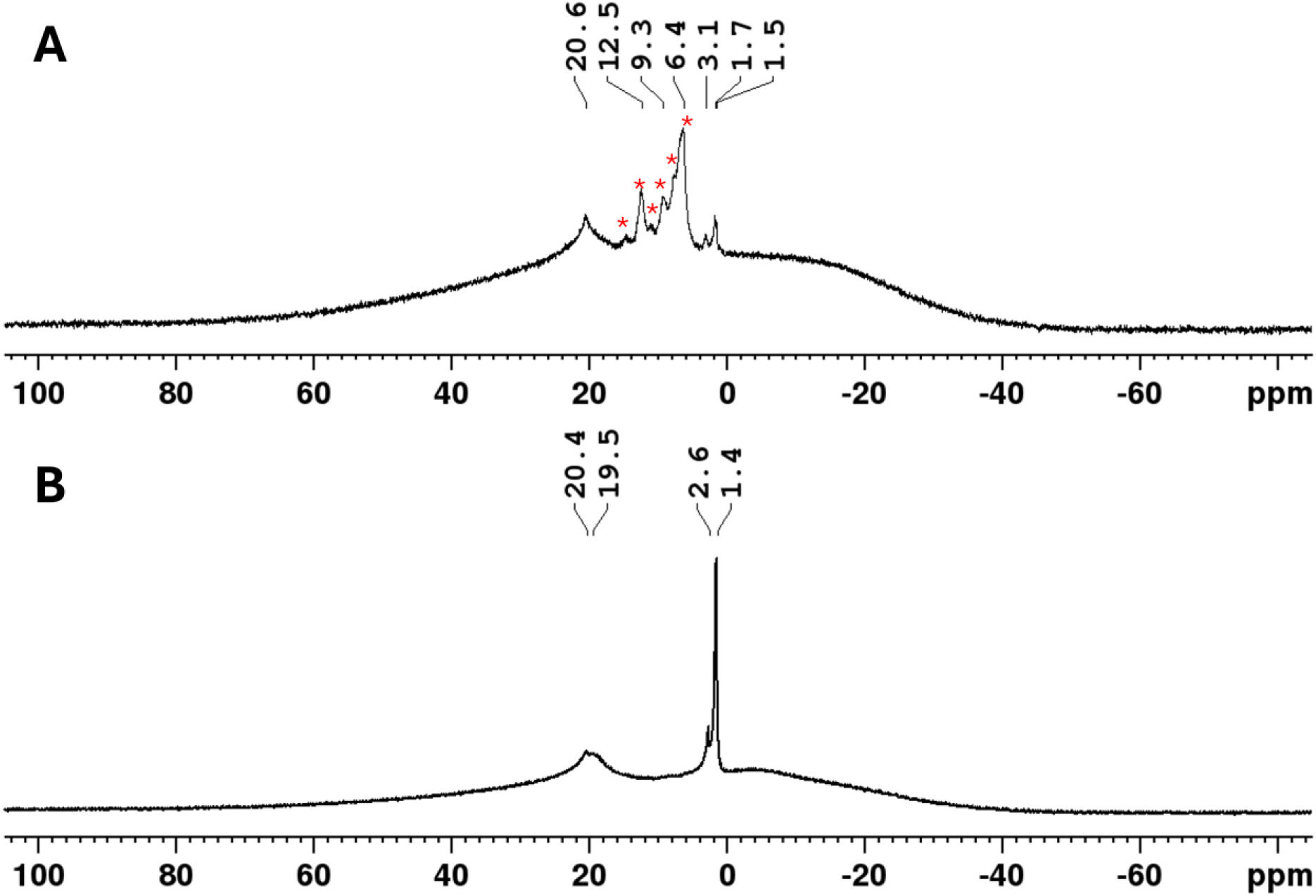
Boric acid reacts with HION to new boron species. A) The ^11^B-NMR spectra showing the crude product of the reaction between HION and boric acid in the presence of a base and B) the crude product of the control experiment performed in the absence of HION. Signals assigned to the new species are marked with an asterisk.

## Discussion

### Identification of novel loci involved in boron homeostasis in maize

Previous analyses of genetic variation in various plant species that detected QTLs or genomic regions associated with boron deficiency or toxicity tolerance, identified mostly boron transporter related genes (Sutton et al., 2007; Zeng et al., 2008; Schnurbusch et al., 2010; Pallotta et al., 2014; Pommerrenig et al., 2018; He et al., 2021; Jia et al., 2021). Notably, the significantly associated SNPs on chr4 and chr7 in the GWAS analysis did not harbor any of the published boron transporter genes in maize, which are located on chr1 (*Zmtls1* and *Zmrte*) and on chr3 (*Zmrte2*) (Chatterjee et al., 2014; Durbak et al., 2014; Leonard et al., 2014; Chatterjee et al., 2017). It is likely that passive, protein-independent boron transport was still prevailing in the low boron field conditions, therefore masking any potential influence of active or facilitated boron transporters. Similar studies using boron concentrations from seed as input have either detected specific boron transporters, like *Zmrte2* (Wu et al., 2021) and uncharacterized genes in maize (Schaefer et al., 2018) or peanut (Zhang et al., 2019), highlighting the potential contribution of non-boron transporter related genes regulating boron homeostasis in plants.

Studies in rice, barley, and peanut have detected various cytochrome P450 proteins (with unknown function or putative iron binding function) to be associated with either boron efficiency traits or boron concentration (de Abreu Neto et al., 2017; Hassan et al., 2010; Zhang et al., 2019), yet *bx* genes, which encode substrate specific cytochrome P450 proteins in grasses, have to our knowledge not been detected as potential candidates influencing boron related processes in plants previously. Our study suggests, that *bx3* represents a novel molecular player associated with boron homeostasis in maize.

While we did not further test the additional GWAS hits detected (Table 1), the identification of a gene encoding a gibberellin receptor GID1L2 (GRMZM2G003246) on chr7 is worth mentioning. A functional role of gibberellin signaling in shaping the maize ionome had been suggested before (Schaefer et al., 2018) and it seems possible that gibberellin signaling or perception might be involved in boron concentration variation. Indeed, a rice GWAS analysis for boron toxicity tolerance also detected a gibberellin receptor (de Abreu Neto et al., 2017), providing evidence for the aforementioned hypothesis and for the applicability of our approach to detect molecular players involved in boron homeostasis. In addition, this finding adds to the importance of phytohormones in the boron response of plants (Martín-Rejano et al., 2011; Abreu et al., 2014; Camacho-Cristóbal et al., 2015; Li et al., 2015; Matthes and Torres-Ruiz, 2016; Eggert and von Wirén, 2017; Poza-Viejo et al., 2018; Gómez-Soto et al., 2019; Zhang et al., 2020; Matthes et al., 2022; Pommerrenig et al., 2022; Matthes et al., 2023) and complements the reports of connections between the adaptation of plants to low or excess boron and gibberellins (Alva et al., 2015; Eggert and von Wirén, 2017).

The unprecedented association between the benzoxazinoid pathway and boron levels is a surprising discovery, particularly because to our knowledge it suggests *bx3* as the first non-boron transporter related gene associated with boron homeostasis in maize. We detected reduced DIMBOA levels and upregulation of various *bx* biosynthesis genes in the boron transporter mutant *Zmtls1* (Table 2), which corroborates previous results showing a transcriptional feedback inhibition of benzoxazinoid biosynthesis genes by DIMBOA (Ahmad et al., 2011). However, altered boron levels and the leaf tip necrosis phenotypes were most pronounced in *bx3* mutants, compared to *bx1* or *bx2* mutants, although all three mutants have reduced DIMBOA levels. Besides a reduction of DIMBOA, the *bx3* mutant, unlike *bx1* or *bx2* mutants, was reported to accumulate the intermediate ION (Abramov et al., 2021). Since the benzoxazinoid pathway is blocked from ION on in the *bx3* mutant, it can be assumed that not just DIMBOA, but also the direct product of BX3, namely HION, is depleted in the *bx3* mutant. Therefore, a depletion of HION or a shift in the ION to HION ratio might be of greater importance for the observed boron-related phenotypes than reduced DIMBOA levels in the *bx3* mutant. This makes it tempting to speculate about a causal connection between these intermediates (ION and/or HION) rather than DIMBOA and enhanced boron levels in the *bx3* mutant. This hypothesis is supported by the finding that transgenic Arabidopsis lines, that overexpress *bx1* and *bx2* and therefore accumulate ION (Abramov et al., 2021), also showed elevated boron levels in rosette leaves and a leaf tip necrosis phenotype (Fig. 5), similar to what was observed in the maize *bx3* mutant (Figs. 1, 2). In the transgenic Arabidopsis lines, ION was further shown to be hydroxylated and glucosylated by endogenous Arabidopsis enzymes to yield a new metabolite, indoline-2-one-5-*β*-D-glucopyranoside (5HIONG; Abramov et al., 2021). Therefore, the Arabidopsis boron-related phenotypes might not just be connected to enhanced ION levels, but also to different intermediates/metabolites, like 5HIONG and/or ratio shifts of ION to such additional metabolites. Furthermore, the detection of additional boron species using ^11^B NMR and ^1^H NMR analyses of HION and boric acid mixtures (Fig. 6), provide compelling evidence for the ability of HION to form a complex with boric acid. While the identity of the additional boron species and proof for such a complex *in planta* remains elusive, it is tempting to speculate that a mechanistic link between boron levels and the benzoxazinoid pathway might be connected to the formation of boron-benzoxazinoid complexes, hence affecting boron homeostasis. While this remains intriguing and speculative the association of boron related processes with the benzoxazinoid pathway opens up exciting new lines of research in both areas.

### Elevated boron levels toxic, beneficial, or both?

The *bx3* mutant, grown in low boron field conditions, showed elevated boron concentrations in the ear leaves, yet boron toxicity phenotypes were subtle at maturity (reduced plant height, increased tillering) (Fig. 1, Supplemental Fig. S5), suggesting that the enhanced boron levels were not high enough to be detrimental to plant performance. On the contrary, strong leaf senescence phenotypes were observed at the seedling stage (Fig. 1H, Fig. 2), indicating that elevated boron levels are more detrimental during juvenile vegetative development, although there are also additional factors that contribute to the leaf senescence phenotype (Fig. 2, Supplemental Fig. S5). Boron requirements are higher during reproductive development compared to vegetative development in maize (Lordkaew et al., 2011; Durbak et al., 2014) and *bx* gene expression is higher during seedling development (Frey et al., 1995; Frey et al., 1997), providing explanations for the leaf senescence phenotype observed in *bx3* seedlings, but not in mature plants. Since boron soil levels at the field site in Bonn-Endenich can be considered low (soil boron concentration: 0.27 mg kg^-1^), it seems possible that the enhanced boron concentration in *bx3* mutants at maturity might counteract soil induced boron deficiency in contrast to provoking toxicity symptoms. This hypothesis is supported by the finding that specific tassel traits, like tassel length and the length of the central spike, but also ear traits, like ear length, were significantly longer in *bx3* compared to B73 plants (Fig. 1G, Supplemental Table S5). This observation further complements published results, showing that boron supplementation increases tassel and ear size (Durbak et al., 2014; Leonard et al., 2014; Matthes et al., 2018). Our findings therefore allow a future assessment of *bx3* and potentially other *bx* genes as suitable molecular candidates for the adaptation of maize to low boron soil conditions.

### Conclusion

Understanding the molecular regulation of boron homeostasis in crops remains a challenging task. By using a GWAS approach in combination with mutant and chemical analyses, we showed that the benzoxazinoid pathway is linked to boron homeostasis likely through *bx3* and the pathway intermediates ION and HION.

## Acknowledgments

We are indebted to the Soil and Plant Testing Laboratory of the University of Missouri, to Angelika Veits and Nur Gömec (University of Bonn – Plant Nutrition) for help with boron measurements. We further thank Chris Browne and his staff, as well as Helmut Rehkopf and Christa Schulz for exceptional plant care during the Missouri and Bonn field seasons, respectively. We are grateful to Laine Weiskopf for assisting in leaf material grinding, to Sherry Flint-Garcia and her team for help with the propagation and providing seeds of the 282 Goodman-Buckler association panel, and to Georg Jander and Kevin Ahern for providing seeds for the *bx2* mutant and the respective W22 inbred control. We thank the Proteomics & Metabolomics Facility (RRID:SCR_021314), Nebraska Center for Biotechnology at the University of Nebraska-Lincoln for the DIMBOA analysis. The facility and instrumentation are supported by the Nebraska Research Initiative.

This work was supported by the Agriculture and Food Research Initiative Grant 2015-06592 from the USDA National Institute of Food and Agriculture to P.M. and by the German Research Foundation (DFG: MA 9520/1-1 and MA 9520/2-1 to MSM). G.S. acknowledges funding by the DFG under Germany’s Excellence Strategy – EXC 2070 – 390732324 (PhenoRob) and A. N.-K. thanks the DFG for an Emmy-Noether Fellowship (NO 1459/1-1) and the Hector Fellow Academy (HFA) for financial support. J.N. thanks the Hector Fellow Academy (HFA) for a Ph.D. scholarship.

The authors declare no conflict of interest.

## Author contributions

V.S.: BLUPs and back-transformed BLUPs generation, GWAS analysis, correlation analysis, data analysis. L.C: Boron concentration measurements in soil, *bx3* mutants and Arabidopsis lines, data analysis, phenotypic analysis. C.S.: Phenotypic analysis of Arabidopsis lines and *bx3* mutants. J.N., T.D.: Boron-benzoxazinoid complex experiments. G. M., Z. D., T. K.: Boron analysis 282 Goodman-Buckler association panel. A.A: Generation and propagation of the Arabidopsis lines used. M.F., A.N.K., H.J., G.S., F.H., P.M., R.A.: Experimental design, student supervision, and data analysis. M.M.: Conceived and designed the study, experimental design, data analysis, student supervision, writing of manuscript with input from all authors. All authors approved of the final version of this manuscript.

## Supplemental information

### Supplemental Material and Methods

Extended Material and Methods

### Supplemental Tables

**Supplemental Table S1:** Boron concentration of maize ear leaves in 277 lines of the 282 Goodman-Buckler association.

**Supplemental Table S2:** Descriptive statistics summary of boron concentrations (µg g^-1^ dw) obtained from the 277 backtransformed BLUPs of the 282 Goodman-Buckler association panel. dw = dry weight

**Supplemental Table S3:** Correlation between gene expression of candidate genes with boron concentration in 277 lines of the 282 Goodman-Buckler association panel.

**Supplemental Table S4:** IPC-OES analysis of various nutrients in B73 and *bx3* mutants.

**Supplemental Table S5:** Phenotypes at maturity of *bx1*, *bx2*, *bx3* mutants and their respective inbred line controls (B73, W22).

**Supplemental Table S6:** Vegetative phenotypes of *bx3* mutants and B73 control plants 14 and 25 days after planting (DAP).

**Supplemental Table S7:** Vegetative phenotypes of *bx3* mutants and B73 control plants 14 days after planting (DAP) with different watering regimes.

**Supplemental Table S8:** Correlation analysis of *bx* gene expression with boron concentration in the 282 Goodman-Buckler association panel.

**Supplemental Table S9:** Phenotypic analysis of *bx1*, *bx2*, *bx3* mutants and their respective inbred line controls (B73, W22) 14 days after planting (DAP), 18 DAP, and 25 DAP.

**Supplemental Table S10:** Leaf necrosis score of Arabidopsis lines, overexpressing parts of the maize benzoxazinoid pathway and *Col-0*.

### Supplemental Figures

**Supplemental Figure S1:** Scoring scheme for assessing leaf necrosis.

**Supplemental Figure S2:** Image segmentation using ImageJ (Schneider et al., 2012).

**Supplemental Figure S3:** Phenotype of the *Zmtls1* mutant grown in the field in Bonn-Endenich or in the greenhouse and boron concentrations of *bx3* mutants.

**Supplemental Figure S4:** Phenotypes induced by boron supplementation and boron concentration in WT and *bx3* ear leaves grown in the greenhouse.

**Supplemental Figure S5:** Leaf necrosis and tassel phenotypes of the *bx3* mutant grown in the field Bonn-Endenich (2020).

**Supplemental Figure S6:** Benzoxazinoid biosynthesis pathway in maize and lactam/lactim tautomerism von HION.

**Supplemental Figure S7:** Boric acid does not react with indolin-2-one (ION) to new boron species.

**Supplemental Figure S8:** Boric acid reacts with HION to form new boron species. A) The ^1^H NMR spectra showing the crude mixture of the reaction of HION with boric acid in the presence of a base, B) the control experiment without boric acid, and C) pure HION.

